# Ancestral and neo-sex chromosomes contribute to population divergence in a dioecious plant

**DOI:** 10.1101/550962

**Authors:** Felix E.G. Beaudry, Spencer C.H. Barrett, Stephen I. Wright

## Abstract

Empirical evidence from several animal groups suggests that sex chromosomes may disproportionately contribute to reproductive isolation. This occurs particularly when sex chromosomes are associated with turnover of sex determination systems resulting from structural rearrangements to the sex chromosomes. We investigated these predictions in the dioecious plant *Rumex hastatulus*, which is comprised of populations of two sex chromosome cytotypes. Using population genomic analyses, we investigated the demographic history of *R. hastatulus* and explored the contributions of ancestral and neo-sex chromosomes to population genetic divergence. Our study revealed that the cytotypes represented genetically divergent populations with evidence for historical but not contemporary gene flow between them. In agreement with classical predictions, we found that the ancestral X chromosome was disproportionately divergent compared with the rest of the genome. Excess differentiation was also observed on the Y chromosome, even when using measures of differentiation that control for differences in effective population size. Our estimates of the timing of the origin of the neo-sex chromosomes in *R. hastatulus* are coincident with cessation of gene flow, suggesting that the chromosomal fusion event that gave rise to the origin of the XYY cytotype may have also been a key driver of reproductive isolation.

## Introduction

Sex chromosomes have long been thought to be associated with speciation (Coyne and Orr 2004). JBS Haldane noticed that when hybrids of one sex were sterile or inviable, it was more often the sex with heteromorphic sex chromosomes (“Haldane’s Rule”; Haldane 1922). A similar, but distinct, observation was later made for X chromosomes. These were found to disproportionately contribute towards hybrid sterility and inviability (the “Large-X” Effect; Dobzhansky 1936; Orr 1989; Coyne 1989) due to a higher density of genes on the X in which introgression had negative fitness effects (Masly and Presgraves 2007). Despite these results, the broad generality of these patterns remains uncertain (Presgraves 2018; Coyne 2018), especially in plants, where dioecy and sex chromosomes have both evolved relatively recently from hermaphroditic ancestors.

Chromosomal rearrangements involving sex chromosomes, including fusions, fissions and translocations, cause evolutionary turnover in sex chromosomes between populations. The newly formed sex chromosomes that arise by this process are referred to as neo-sex chromosomes (White 1940). Fusions and translocations involving sex chromosomes may increase the influence of sex chromosomes on the divergence process (Connallon et al. 2018), and rearrangements have been implicated in driving divergence between populations (White 1978; Rieseberg 2001; Guerrero and Kirkpatrick 2014; Ortiz-Barrientos et al. 2016). Furthermore, sex chromosome turnover can cause imbalance in the sex determination of hybrids resulting in hybrid infertility and divergence between populations with different sex chromosome systems (Graves 2016). For these reasons, sex chromosome turnover is thought to be a particularly important driver of population divergence and speciation.

Speciation has often been investigated through crosses between divergent populations and assessment of their interfertility; however, recent advances in sequencing technology offer population genomic approaches as a complementary, albeit indirect, avenue for exploring patterns of genetic divergence. Genome-wide differences in allele frequencies and the fixation of alternate alleles between populations can provide insights on the extent of divergence and gene flow (Wang et al. 1997). By comparing genetic diversity and divergence on X chromosomes, Y chromosomes, neo-sex chromosomes and autosomes, it is possible to infer whether sex chromosomes have played a disproportionate role in the evolution of reproductive isolation. Using such approaches, sex chromosome turnover has been implicated in population divergence and speciation using genomic data from stickleback (Kitano et al. 2009), swordtail (Franchini et al. 2018), pine-beetle (Bracewell et al. 2017), and butterfly (Smith et al. 2016). However, the role of sex chromosome turnover in the genetic divergence of populations and incipient speciation processes has not been investigated in plants.

To investigate the patterns of divergence on plant sex chromosomes, we measured population-wide genetic variation in the dioecious, wind-pollinated, annual flowering plant *Rumex hastatulus*. This species is polymorphic for sex chromosome karyotype: individuals in populations that are distributed west of the Mississippi river, USA have four sets of autosomes, males with X and Y chromosomes and females with a pair of relatively small X chromosomes. In contrast, individuals in populations to the east of the Mississippi have three autosomes, males with an X and two Y chromosomes and females with a pair of large X chromosomes (Smith 1964). Cytological evidence suggests that these karyotypic differences can be explained by a reciprocal translocation between an autosome and the X chromosome in an ancestor with the XY karyotype resulting in a cytologically larger X and an extra Y chromosome in descendants (Smith 1964). A recent cytological study using florescent probes on hybrid cytotypes has provided evidence consistent with this scenario (Kasjaniuk et al. 2018). The two geographical groups in *R. hastaulus* are hereafter referred to as cytotypes (and are akin to the groups referred to as “races” in Smith, 1964). Investigations of synthetic polyploids indicate that the cytotypes have different modes of sex determination; for the XY cytotype, maleness is determined by presence of the Y chromosome, whereas in the XYY cytotype, maleness is determined by the X to autosome ratio (Bartkowiak 1967). Such karyotype changes in the sex chromosomes enable tests of the effects of rearrangements and changes in sex determination on population genetic divergence, an approach we use here.

Several earlier findings are consistent with the hypothesis that the two cytotypes of *R. hastatulus* are at least partially reproductively isolated and may represent early stages in the speciation process. The different cytotypes are differentiated in phenotypic morpho-space (Jackson 1967; Simpson 2013) and recent crossing experiments found evidence of sex-specific asymmetrical reduced viability of cytotype hybrids (Kasjaniuk et al. 2018). However, the timing of divergence and extent of historical gene flow between the cytotypes in nature remains unclear. Previous estimates of divergence between the cytotypes based on 13 nuclear loci estimated divergence at less than 150,000 years ago (Simpson 2013), whereas analysis of transposable element accumulation patterns estimated divergence times of roughly 600,000 years ago (del Bosque et al. 2011). Model-based coalescent approaches for inferring the timing of divergence and extent of gene flow between the cytotypes have not been conducted and are therefore a focus of this study. These approaches are particularly important when trying to infer the potential role of sex chromosomes in population divergence and reproductive isolation. This is because it is only under models of ’complex speciation’ with gene flow during divergence that we expect a potential role for sex chromosomes in reproductive barriers (Presgraves 2018). Here, we build the first comprehensive evolutionary demographic model of *Rumex hastatulus* and use population genomic approaches to investigate whether ancestral or neo-sex chromosomes contribute disproportionately to reproductive isolation in comparison to genes residing elsewhere in the genome.

## Methods

### Sample collection, DNA extraction and sequencing

*Rumex hastatulus* RNAseq population samples were previously sequenced (Hough et al. 2014) and we supplemented that data set with reduced representation sequencing for 93 more individuals of which 23 males were also sequenced using RNAseq. Information on sex ratios and demographic data for all of the populations we examined is available in Pickup and Barrett (2013) and additional sampling information for the individuals used in this study is summarized in Table S1.

We extracted DNA from leaf tissue of 93 individuals and obtained genomic extractions using the DNeasy Plant Mini Kit (QIAGEN GmbH, Hilden, GERMANY) according to manufacturer’s protocol. Genomic DNA was sequenced using Genotyping-By-Sequencing (Elshire et al. 2011). For producing the reduced representation library, we digested genomic DNA using the restriction enzyme PstI, and libraries were sequenced at the Australian Genome Research Facility (AGRF). For a subset of 23 males, we also collected RNA from leaf tissue. We performed RNA extractions using Spectrum Plant 159 Total RNA kits and the RNA was sequenced using a poly(A) mRNA library with prep and sequencing performed at The Center for Applied Genomics in Toronto, ON.

### Alignment and SNP calling

After sequencing, we removed low quality reads from further analysis using an in-house script (posted to https://github.com/cafeblue/wei_script/blob/master/clean_B_N_HiSeq.pl; last updated December 6, 2017): briefly, reads with 50% of bases with a quality score <23, or with <10% of non-ambiguous bases were classified as low quality and removed. We mapped all *R. hastatulus* reads to the transcriptome assembled in Hough et al. (2014), where predicted transcripts were subsequently trimmed to be in reading frame. The GBS and RNAseq files were analysed independently. Mapping was conducted as previously described (Beaudry et al. 2017), which generally follows GATK best practices (McKenna et al. 2010). We produced primary alignments with BWA mem (Li and Durbin 2009), allowing multiple mapping but not discordant mapping, and using default conditions: a mismatch penalty score of 4, a minimum seed length of 19, a band-width of 100, and a gap penalty of 6. We remapped the primary alignment using STAMPY (Earl and vonHoldt 2012) under default parameters, with the “–bamkeepgoodreads” option on, a substitution rate of 0.001, and disallowing either multiple mapping or discordant mapping. We processed mapping files using PICARD’s AddOrReplaceReadGroups followed by MarkDuplicates, with default parameters (https://broadinstitute.github.io/picard/; last accessed December 6, 2017). We called SNPs for each individual using GATK (McKenna et al. 2010) Haplotypecaller with the -ERC GVCF option, and merged the g.vcf files using GATK GenotypeGVCFs. We filtered polymorphisms using GATK VariantFiltration with a cutoff site quality score of >50 and retained only biallelic SNPs using GATK SelectVariants.

### Demographic analyses

We used clustering-based methods to investigate genetic differentiation between the *R. hastatulus* cytotypes. For the first round of demographic analyses, both RNAseq and GBS datasets were analyzed, with a focus on the GBS set as it represented greater within-population sampling. We began this series of analyses by examining genetic structure by randomly subsampling SNPs with STRUCTURE v2.3.4 (Pritchard et al. 2000). STRUCTURE was run with a 1,000 burnin, for a subsequent 10,000 generations. We tested K-values of 1-7 and ran five replicates for each value of K. We determined the most likely K from likelihood scores using StructureHarvester (Earl and vonHoldt 2012), according to the method of Evanno et al. (2005). We used CLUMPP (Jakobsson and Rosenberg 2007) permutations to align clusters across runs and plotted results using *distruct* (Rosenberg 2004). We also used genotype calls to create neighbour-joining trees with *ape* (Paradis 2011) in R and used splitstree (Huson 1998) to visualize recombination events between populations. To further test for genetic structure, we employed Principal Component Analysis (PCA) with FactoMineR (Husson et al. 2010) in R.

Clustering of populations in such analyses as those performed above can be caused by geographic discontinuity in sampling (Balkenhol et al. 2015; Bradburd et al. 2016; Perez et al. 2018). In such cases, clustering may be a product of the interaction between incomplete sampling and isolation-by-distance rather than discrete barriers to gene flow. As our sampling did not include the purported cytotype contact zone (Smith 1964), and thus creates a discontinuity in sampling, our results may be influenced by sampling design. This is of particular concern here as our PCA suggested isolation-by-distance played a significant role in structuring genetic variation within each *R. hastatulus* cytotype. To evaluate this possibility, we examined the Estimated Effective Migration Surface (EEMS) (Petkova et al. 2016), which is a Bayesian estimate of effective migration rates based on SNP frequencies and a prior hypothesis of isolation-by-distance. Effective migration here refers to how often alleles successfully spread and not to potential rates of dispersal. To use EEMS, we converted the .vcf of filtered SNPs to .plink using vcftools, and, as we did not impute data, we used bed2diff_v1 (in the EEMS package) to convert .bed files from plink to the EEMS format. We ran six MCMC chains through the EEMS pipeline, before exporting into EEMS package in R for synthesis of MCMC results and subsequently for visualization. Observed and fitted dissimilarities revealed no outliers (Fig. S1).

To further assess signals of gene flow, we used the joint allele frequency spectrum to test various demographic models with the software δaδi (Gutenkunst et al. 2009). We used the gene annotation from fgenesh as assessed in Crowson et al. (2017) to find synonymous SNPs in our RNA dataset. We extracted synonymous SNPs from the annotated VCF using SNPeff (Cingolani et al. 2012) and these were filtered for a quality score higher than 50, and a coverage depth higher than 5 for every individual. We converted the synonymous SNP VCF to δaδi input using vcf2dadi.pl (https://github.com/owensgl/reformat/blob/master/vcf2dadi.pl; last access April 5 2018). In δaδi, we subsampled individuals of the XY and XYY cytotypes to a projection of 38 and 48 alleles, respectively. The partial differential equation approximating the frequency spectrum was solved at phi-array = [60,70,80], before extrapolation to an infinitely fine grid. We tested 21 models from dportik (https://github.com/dportik/dadi_pipeline; last access: Dec 18, 2018) (Portik et al. 2017). We chose the best model using AIC and confidence intervals were calculated using the Goodness of Fit pipeline in dportik dadi_pipeline: we resampled in 3 rounds of 20, 30 and 50 reps of optimization respectively, over 100 simulations in each round. We used Polymorphurama (Bachtrog and Andolfatto 2006) to get an estimate of the total number of synonymous sites in our transcriptome assembly. We divided δaδi’s estimates of Θ by 1,288,938 total synonymous sites. We roughly estimated the effective population size (*N*e) as Θ/4Lµ, using the *Arabidopsis thaliana* estimate of the mutation rate of 7 x 10^-9^ base substitutions per site per generation (Ossowski et al. 2010).

At neutral sites, substitutions between populations should accumulate at the rate of incoming mutations (2µ; Kimura, 1983). This prediction from the neutral theory suggests that, given the mutation rate, time since divergence can be estimated from the amount of divergence between two populations (Li 1997). The neutral rate of divergence technique has been used to estimate the time since different strata on sex chromosomes have stopped recombining (Lahn and Page 1999). This method assumes no further genomic exchange such as recombination between the sex chromosomes and that the expected pairwise coalescence time on either sex chromosome is the same now as it was for the autosomal ancestor. Using this technique, we calculated the time since recombination stopped between the neo-sex chromosomes. Previous work estimated the mean K_S_ of neo-X branches as 0.00276 and of the neo-Y branch as 0.00297 (Hough et al. 2014). To control for ancestral polymorphism, which should be a component of divergence between species, we subtracted the mean π_syn_ value of the two cytotypes for autosomal loci from K_S_, before dividing by 2µ, to give our estimate of time since divergence (Hudson et al. 1987).

### Diversity and differentiation estimates

To investigate genome-heterogeneity in diversity and differentiation, genes were identified as sex-linked, neo-sex-linked following Hough et al. (2014) and as autosomal according to Hough et al. (2017). Sex-linked SNPs were phased, using methods described below. For the other sets of loci, we called SNPs for each individual using GATK (McKenna et al. 2010) UnifiedGenotyper, and .vcf files were converted to .fasta using vcf2fasta_uni.pl (by Wei Wang), according to the details of fasta_maker_v2.sh which also removes sequences with less than 60% coverage. Scripts are available at felix.beaudry’s GitHub reads2poly repository (https://github.com/felixbeaudry/reads2poly; Feb. 10, 2019 update) .

We phased genes identified as sex-linked (see above) using HapCut (Bansal and Bafna 2008). Haplotypes were identified as X or Y depending on sex-specific SNPs, following Hough et al. (2017). Briefly, reference alleles heterozygous in all males and homozygous in all females were identified as X-linked, while the non-reference allele, absent in all females, was identified as Y-linked. During this analysis, we discovered that three individuals with male phenotypes (populations NCELI, NCHIC and FLHAM) had none of the male-specific SNPs. As inconstant males have been recorded previously in *R. hastatulus* (for example, Smith 1963), these samples were removed from all sex chromosome analyses. As hemizygous genes may have significantly different dynamics from X loci that still retain their Y homologs (Charlesworth et al. 1987), and also represent a very small subset of X-linked loci for polymorphism estimation, we removed hemizygous genes, as identified in Hough et al. (2014), from further analyses.

For the neo-sex chromosomes, there were too few fixed differences to reliably phase our data between X and Y haplotypes in males. We therefore estimated between-population divergence using only female individuals, as these data will include only neo-X data and therefore avoid the need for phasing. To ensure that this smaller dataset did not influence values of divergence, we compared male and female divergence at other loci and found that there was no significant difference between males and females in divergence for any gene set (Fig. S2). We processed .fasta files using polymorphurama_interpop.pl, edited from Polymorphurama (Bachtrog and Andolfatto 2006).

If sex chromosomes disproportionately contribute to divergence, we expect to see significantly different allele frequency differentiation (for example *F*_ST_ values) at these loci compared to autosomal loci. High *F*_ST_ values can also be caused by low levels of within-population polymorphism, in which case the values may not reflect increased divergence between populations at those loci (Charlesworth et al. 1997). For example, regions of lower recombination are disproportionately affected by linked selection, have lower levels of within population variation, and therefore exhibit high values of *F*_ST_ (Cruickshank and Hahn 2014). To test for this, we complemented our analysis using *F*_ST_ values with Nei’s *d*_XY_ (Nei 1987), which measures the average divergence between pairs of haplotypes from each population and is not affected by changes in current effective population size. However, *d*_XY_ is affected by ancestral effective population size (Cruickshank and Hahn 2014). To remove the effects of ancestral polymorphism on the estimate of divergence, we used absolute divergence, *d*_A_ (Nei 1987), which subtracts the mean value of π between clades from *d*_XY_.

### Testing for introgression

To investigate the possibility of Y introgression, we concatenated X and Y phased haplotypes separately, only using loci where coverage was present in all individuals. We constructed Maximum Likelihood trees in RaxML v8.2.10 (Stamatakis 2014), using a GTR-GAMMA model. The best tree was chosen from 20 trees and bootstrapped 100 times. To assess the likely timing of divergence between the Y haplotype clusters, we approximated neutral divergence by subtracting mean π_syn_ for each Y haplotype cluster from the K_S_ value between each Y haplotype cluster and divided by 2µ. Both values of K_S_ and of π_syn_ were estimated using Polymorphurama.

To test for genome-wide introgression, we performed an ABBA-BABA test (Green et al. 2010; Patterson et al. 2012) using the evobiR package (Blackmon 2016) in R on the set of autosomal loci. We tested for excess allele sharing between the XY cytotype and individuals from the Southern range of the XYY cytotype using the top BLAST hit from the *Rumex rothschildianus* transcriptome assembly (Crowson et al. 2017) as an outgroup sequence. To estimate the standard deviation of Patterson’s D, we performed 1000 jackknife replicates with a block size of 1000 SNPs.

## Results

### Genetic differentiation of the cytotypes

After filtering for coverage, quality and missing data, the GBS set has 5,970 SNPs and the RNAseq set 916,255 SNPs. We found there were no sites with fixed differences between cytotypes in the GBS dataset and only 136 in the RNAseq dataset (Table 1). The majority of SNPs were uniquely segregating within one or the other cytotype, although 14% of GBS SNPs and 21% of RNAseq SNPs were shared between the two groups.

**Table 1.** Number of shared polymorphisms and fixed differences between the *Rumex hastatulus* cytotypes

|  | GBS | RNA |
| --- | --- | --- |
| Fixed SNP differences | 0 | 136 |
| Shared SNPs | 672 | 191,506 |
| XY-unique SNPs | 1,879 | 387,692 |
| XYY-unique SNPs | 2,243 | 330,141 |

Using STRUCTURE, we identified K=2 as the most likely number of population clusters in our GBS dataset (Fig. S3A). As expected, individuals clustered according to cytotype (Fig. 1A, B). Visualization at other values of K did not reveal further structure between the cytotypes (Fig. S3D), and there was no clear evidence for recent introgression between the groups. Splitstree also clustered populations by cytotype (Fig. S4), and the same pattern of grouping held using the RNAseq data (Fig. S5). Within each of the XY and the XYY cytotypes, we found K=3 and K=2, respectively, as the most likely number of clusters using STRUCTURE. However, visualization suggested that these clusters do not represent within-cytotype substructure (Fig. 1C, D).

**Figure 1.**
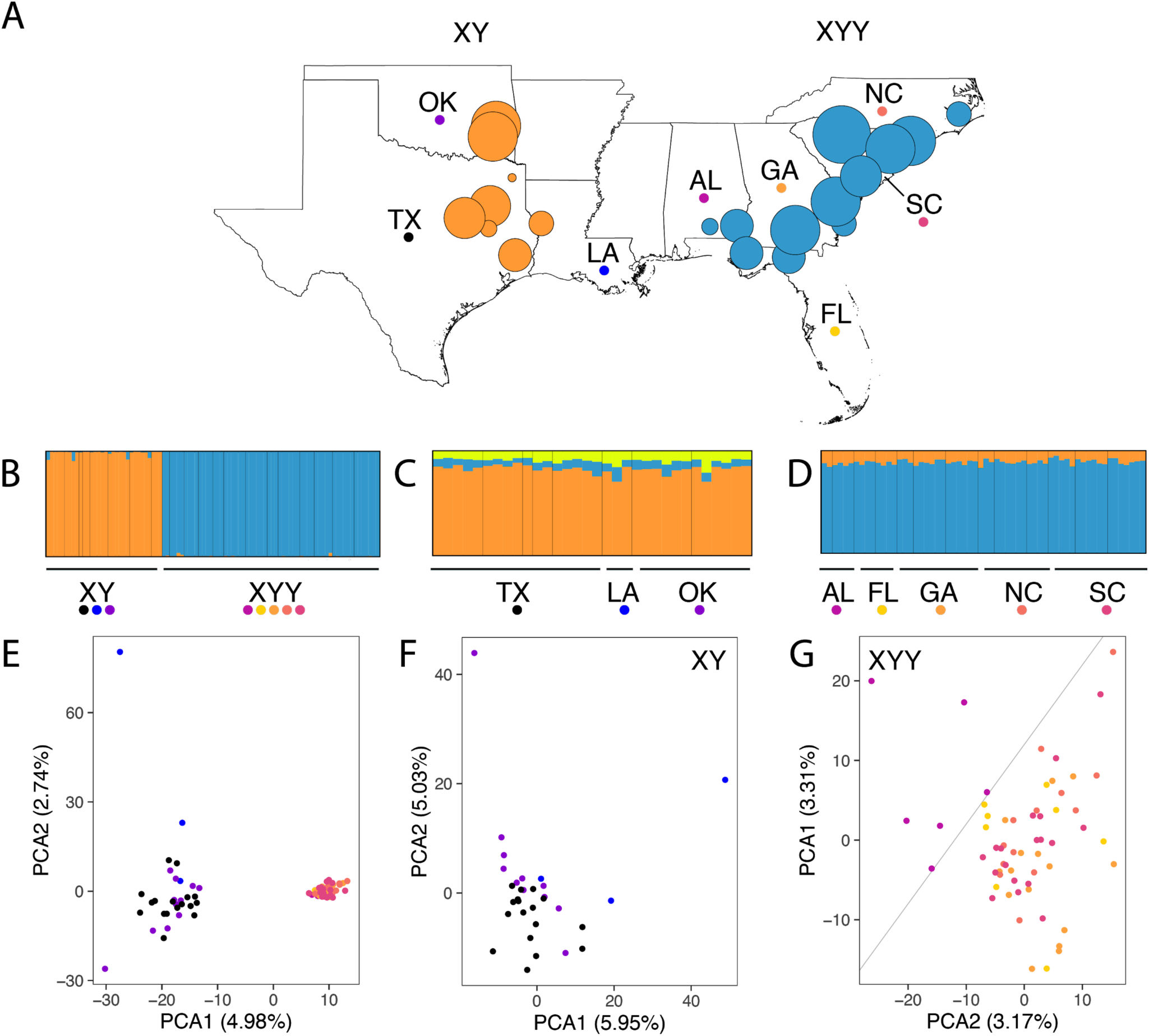
**A,** Geographical sampling of *Rumex hastatulus* populations used for the Genotype-by-Sequencing (GBS) dataset, colored according to cluster predicted from the STRUCTURE analysis; **B-D,** STRUCTURE plot for individuals (**B)** across the range of *R. hastatulus*, (**C**) from the XY cytotype, and (**D**) from the XYY cytotype, labelled by state of provenance **E-G**, Principal Component Analysis (PCA) for individuals (**E)** across the range of *R. hastatulus*, (**F**) of the XY cytotype, and (**G**) of the XYY cytotype

To further investigate population structure of *R. hastatulus*, we perform Principal Component Analysis (PCA) for both cytotypes together, as well as for each separately. When considering all samples, the results of the PCA replicated the pattern of clustering found using STRUCTURE (Fig. 1E). PC1, which explained 4.98% of the variance, clearly separated populations of the XY cytotype from those of the XYY cytotype. Within each cytotype, some of the variation along the PC axes can be explained by isolation-by-distance. In the XY cytotype, PC1 correlates with longitude (East-West axis; linear model, *R*^2^ = 0.18, *P* = 0.016) and PC2 shows a nonsignificant trend towards a correlation with latitude (North-South axis; linear model, *R*^2^ = 0.11, *P* = 0.069; Fig. 1F). In the XYY cytotype, PC2 correlates with longitude (East-West axis; linear model, *R*^2^ = 0.24, *P* = 8.63e-05; Fig. 1G), but the relation between PC1 and latitude is not significant. Both PC1 and PC2 suggest Alabama populations are somewhat distinct from other XYY populations (gray line in Fig. 1G), so the relation between PC1 and latitude in XYY populations may be complicated by the effect of additional divergence in populations from Alabama (AL). Once populations from Alabama were removed, PC1 showed a nonsignificant trend towards a correlation with latitude (linear model, *R*^2^ = 0.0731, *P* = 0.06). To evaluate the possibility that clustering is due to geographic discontinuity in sampling, we considered the Estimated Effective Migration Surface (EEMS). As predicted, effective migration rates were estimated to be low across the contact zone (warm colors in Fig. 2).

**Figure 2.**
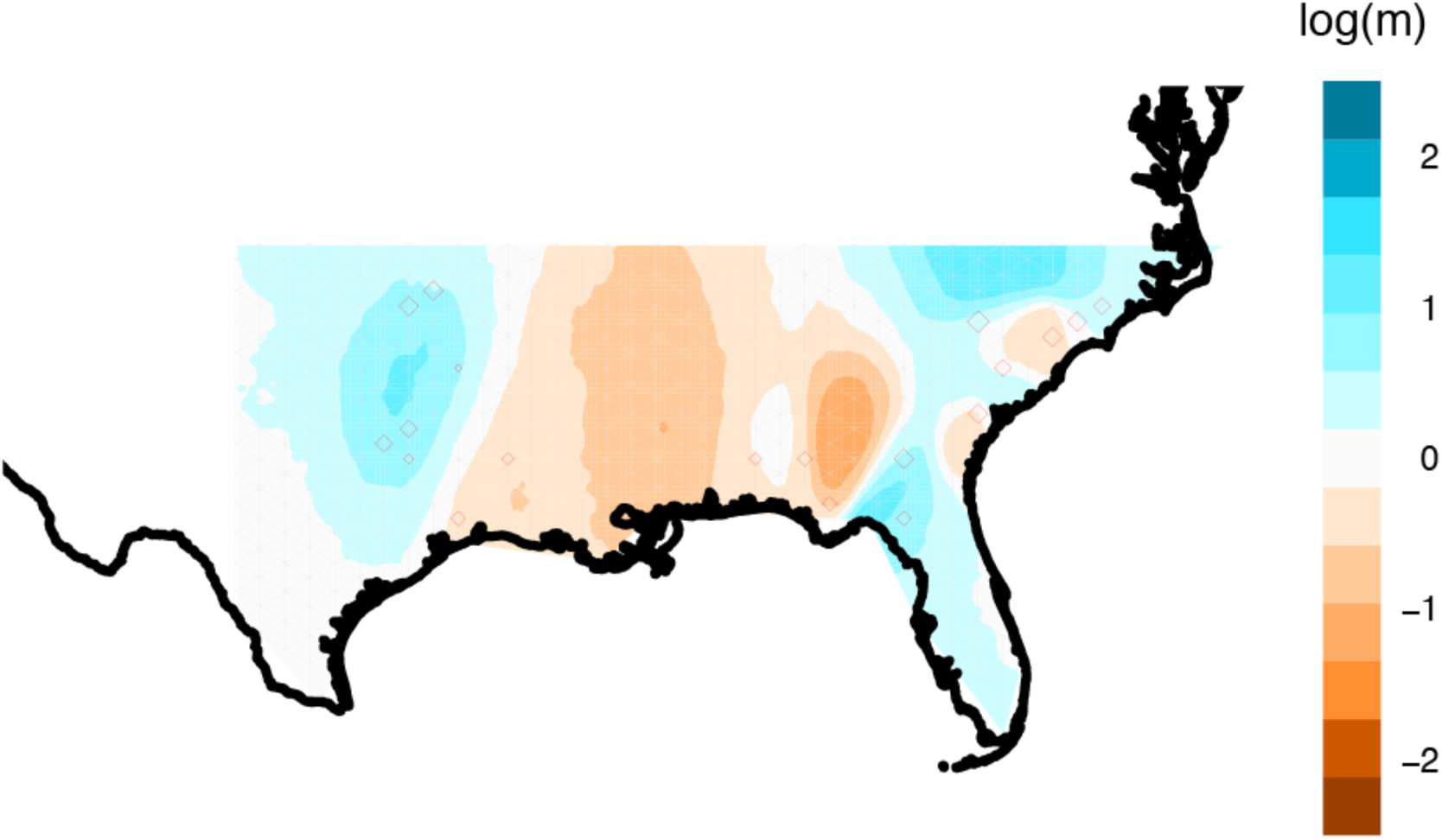
Estimated Effective Migration Surface (EEMS) across the range of *Rumex hastatulus*, colored according to estimated rate of migration: cold colors represent high levels of migration and warm colors represent low levels of migration.

Given these patterns, we sought to distinguish whether reproductive isolation has evolved in strict allopatry, or whether there has been a history of ongoing gene flow concurrent with genetic divergence. To investigate this, we created and fit models of demographic history to autosomal synonymous SNP frequencies from our RNAseq dataset using δaδi. Out of 21 models, our most likely model was one of divergence across two stages: the first and oldest stage showed lowered and asymmetric gene flow, while the second showed no gene flow, and each stage was characterized by different population sizes (lnL = -12668.91, AIC = 25353.82; results summarized in Table S2). This model was significantly more likely than a model of divergence without gene flow (LRT, d.f. = 5, *P* << 0.001). Given that our previous analyses suggested a unique pattern of divergence in Alabama populations, we reran the model fitting without individuals from Alabama. We found that a model of divergence with ancient asymmetrical migration and size change was again the most likely model, with a better fit to the data than the model including Alabama populations (lnL = -12013.95, AIC = 24043.9; Fig. 3, Fig. S6, results summarized in Table S2). Both models are detailed in the supplementary materials, but for subsequent analyses we only consider the model fit from the data that excluded Alabama populations; however, conclusions drawn are generally consistent between both demographic models.

**Figure 3.**
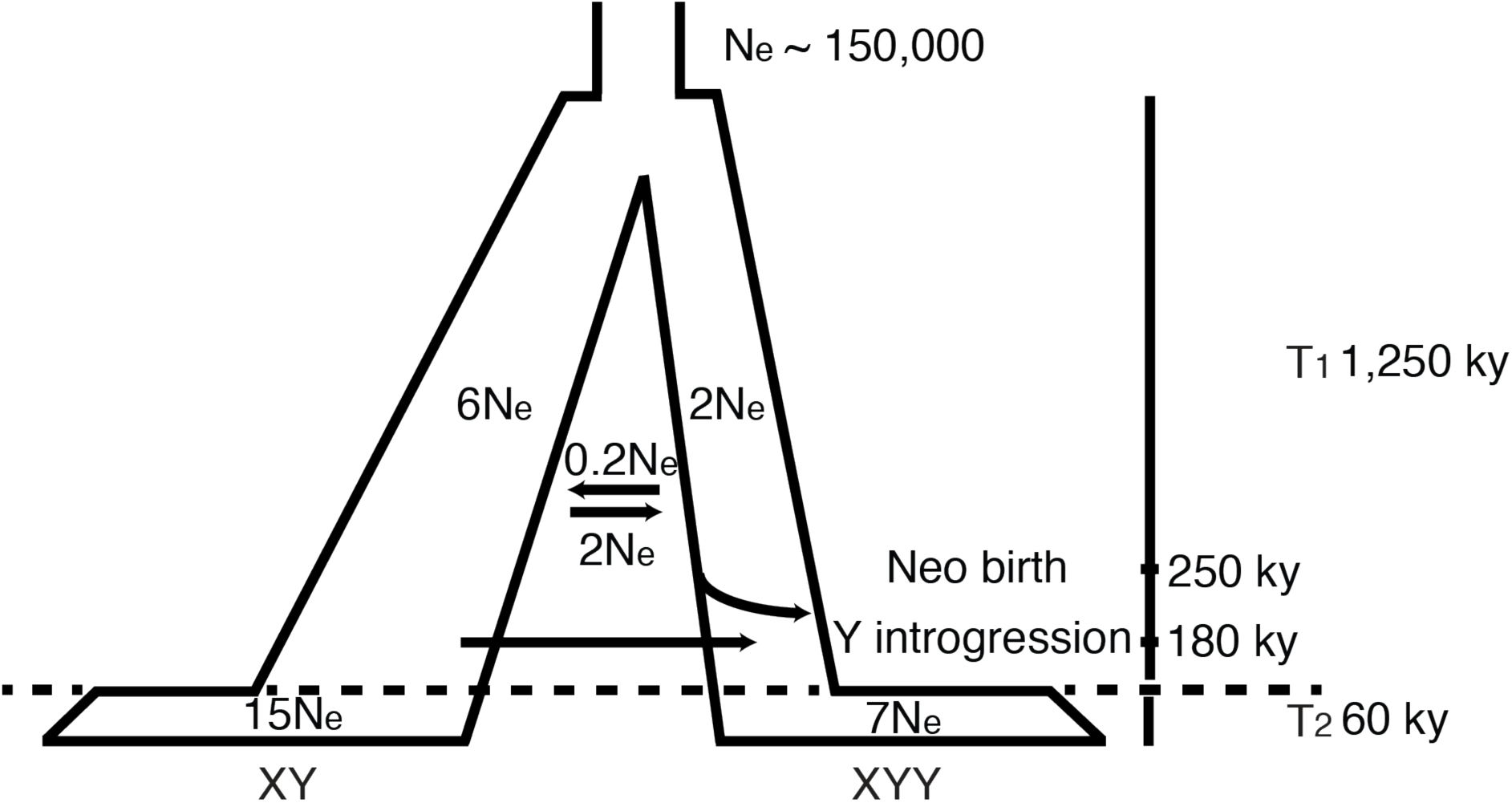
Graphical recreation of the most likely demographic model of *R. hastatulus* from analysis in δaδi. Units are scaled by effective population size (*N*e) or thousands of years (ky).

Under this model, both the XY and XYY cytotypes are inferred to have undergone population expansion upon divergence, with the inferred expansion in the XY cytotype much larger than that of the XYY cytotype. Estimates of migration are asymmetrical between the cytotypes; gene flow from the XY to the XYY is an order of magnitude larger than in the opposite direction. This difference in gene flow estimates cannot be entirely explained by the difference in effective population size between the cytotypes. The model inferred a second stage of divergence 6*N*e generations after initial divergence, characterized by a complete absence of gene flow between the cytotypes, at which time both cytotypes again experienced population expansion.

Using the per base pair nucleotide mutation rate estimate from *Arabidopsis thaliana* and the model’s estimate for T, we estimated that the ancestral *N*e for *R. hastatulus* was approximately 150,000 individuals. With this estimate, we can also estimate time since divergence. The first stage of divergence is estimated at over 1 million years ago, assuming 1 generation per year, given that *R. hastatulus* has an annual life cycle. The start of the second stage of divergence is approximately 60,000 years ago.

To investigate how the timing of divergence relates to the evolution of the sex chromosome systems, we sought to estimate the timing of the sex chromosome rearrangement event. Assuming a neutral rate of divergence, we estimated that the loss of recombination between the neo-sex chromosomes occurred approximately 251,000 years ago. Thus, the neo-sex chromosomes formed prior to our estimate of the timing of complete cessation of gene flow.

### Estimates of divergence on the sex chromosomes

We next sought to further explore whether the sex chromosomes or neo-sex chromosomes disproportionately contribute to divergence in *R. hastatulus*. We first investigated whether there were differences between the X, Y, and neo-sex chromosomes compared to autosomal loci in the variance in allele frequencies between populations, as measured by Wright’s *F*_ST_ at synonymous sites. Consistent with this hypothesis, we found significantly higher *F*_ST_ values at X (Welch Two Sample *t*-test, *P* < 2.15e-05), neo-X (*P* < 0.013) and Y loci (*P* < 7.52e-24) than at autosomal loci (Fig. 4A).

**Figure 4.**
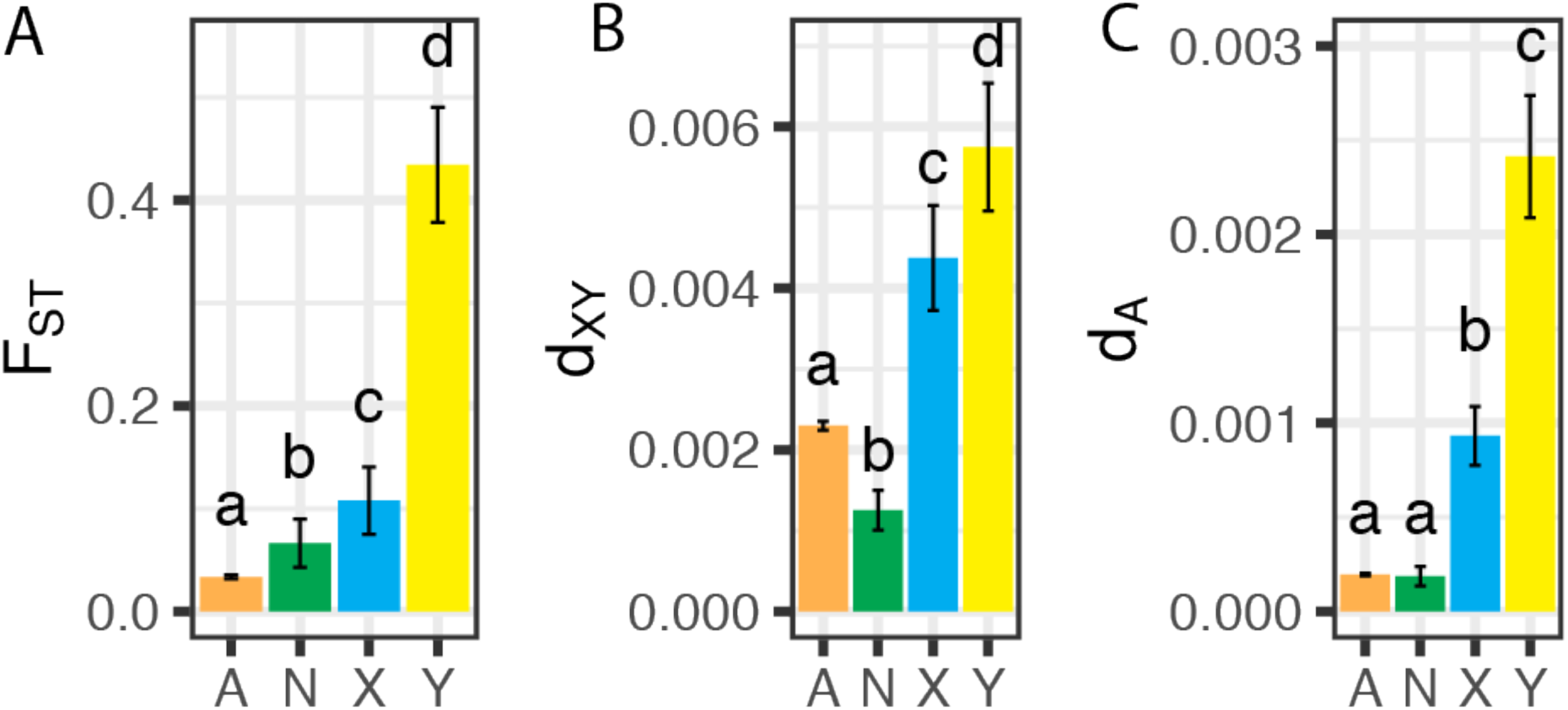
Mean values of **A.** *F*_ST_, **B.** *d*_XY_, and **C.** d_A_ for autosomes (orange), neo-sex chromosome (green), X haplotypes (blue) and Y haplotypes (yellow) in *Rumex hastatulus*.

To test whether high *F*_ST_ values were caused by low levels of within-population polymorphism, we used Nei’s *d*_XY_. Consistent with the prediction of excess divergence at sex chromosomes, we found significantly lower *d*_XY_ values at autosomes than at the X (*P* < 3.77e-10) or Y (*P* < 5.97e-15) chromosomes (Fig. 4B). In contrast, neo-X genes showed significantly lower *d*_XY_ than any other category (*P* < 1.25e-11 compared to autosomal genes). As *d*_XY_ is affected by ancestral polymorphism, we estimated absolute values of divergence, *d*_A_. Using *d*_A_, we found no significant difference between autosomal loci and neo-sex-linked loci (*P* = 0.94), whereas *d*_A_ for the old X and Y chromosomes remained high (Fig. 4C). The pattern holds using pairwise *d*_SYN_ (*P-value* for all pairwise comparison between chromosomes < 1e-6, except for between X and Y where *P-value* = 0.77).

### Polymorphism suggests unidirectional gene flow of Y haplotypes

Linked selection reduces within-population polymorphism, a pattern that should be extreme on a large non-recombining region such as the Y chromosome. Given this expectation, we found a surprising result in the XYY cytotype; π_syn_ at Y loci within the XYY cytotype is similar to π_syn_ at X loci (Fig. S7).

To evaluate whether we can explain this finding by cryptic population structure for Y haplotypes in the XYY cytotype, we constructed a Maximum Likelihood tree of phased X and Y haplotypes from both cytotypes (Fig. S8). X haplotypes were distributed among three clades: 1) X haplotypes from the XY cytotype, 2) most X haplotypes from the XYY cytotype, and 3) a clade sister to XYY-X haplotypes which consists of X haplotypes from Alabama (Figure S8A). In contrast, Y haplotypes were distributed among four clades. Contrary to expectation, haplotypes from the western range of the XYY-populations clustered with Y haplotypes from XY-populations, to the exclusion of Y haplotypes from the Northern range of the XYY cytotype (Fig. 5). Again, Alabama haplotypes formed their own clade, but this subclade fell squarely within the Northern-XYY Y haplotype clade (Fig. S8B). When we estimated genetic diversity in Y haplotypes partitioned by their subclades, we observed the expected reduction in π_syn_ at phased Y loci (Fig. S7A). Nonetheless, values of *F*_ST_ and *d*_A_ between the subclades, which are affected by π_syn_, were significantly higher on the X and Y as compared to autosomal and neo-sex-linked loci (Fig. S7B).

**Figure 5.**
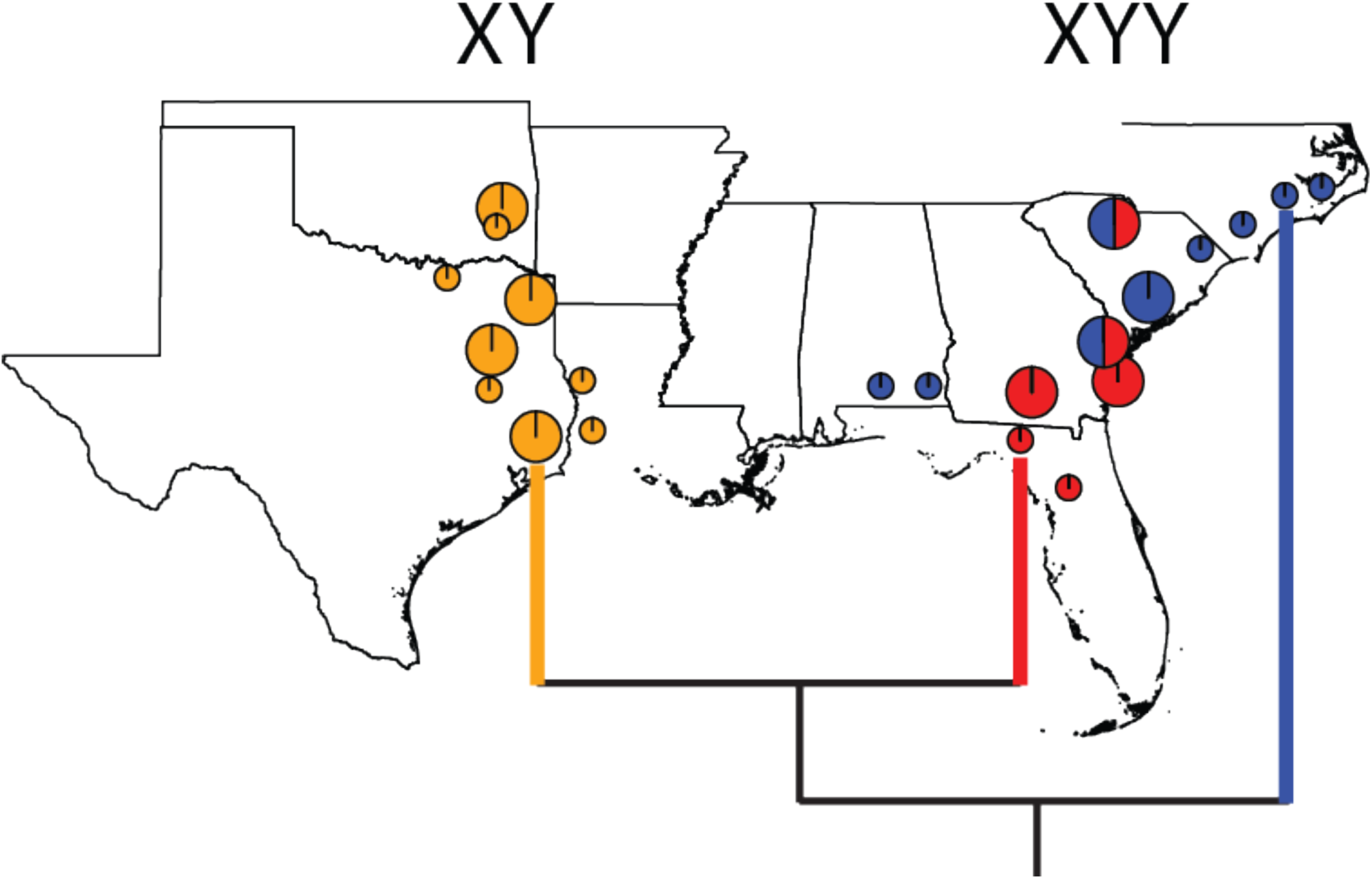
The ML tree of Y haplotypes showing clustering in three clades overlayed on the geographic sampling locations of individual populations of *Rumex hastatulus*.

To investigate whether the pattern of Y haplotype clustering could be explained by gene flow, we tested for a more general pattern of disproportionate gene flow between the XY cytotype and the Southern XYY clade. We ran an ABBA-BABA test on the set of autosomal loci. We found significant evidence for gene flow between these two population samples (D = 0.57 ± 0.0038, Z = 148.65, *P* << 0.001). Given incomplete lineage storing is unlikely to explain Y clustering (Pease and Hahn, 2013), we estimated the timing of XY→Southern XYY Y haplotype introgression. Assuming a neutral rate of divergence, we estimated this event took place roughly 180,000 years ago. Because our estimate for the time since divergence between these Y haplotypes predates the loss of gene flow between the cytotypes, the pattern of Y clustering is more accurately described as resulting from gene flow than introgression.

## Discussion

The history of gene flow between the cytotypes of *Rumex hastatulus* sheds light on the potential role of sex chromosomes in population differentiation. We found evidence for a two-staged loss of gene flow between cytotypes, in which each ancestral and neo-sex chromosome appears to have played a role. Our analyses are consistent with growing evidence in numerous systems that population divergence proceeds with on-going gene flow (Presgraves 2018); this ’complex speciation’ model implies that it is possible for sex chromosomes to contribute disproportionately to reproductive isolation throughout the process of divergence. As we discuss further below, a key open question concerns the relative contributions of geographic isolation, the older sex chromosomes, and the formation of the neo-sex chromosomes in causing reproductive isolation between populations.

### General pattern of divergence and population history

PCA and EEMS analyses suggested that there are two barriers contributing to genetic differentiation, the most important one separating the two cytotypes, and a second separating Alabama populations from the rest of the XYY clade. These results are consistent with geographic barriers, as both the Mississippi river and the Appalachian Mountains have been identified as regions that contribute to reduced gene flow and often participate in enabling incipient speciation in both animal and plant species (Jaramillo-Correa et al. 2009; Soltis et al. 2006; Pyron and Burbrink 2010).

Our inference of demographic history implies a complete loss of gene flow between the two cytotypes within the last 60,000 years. However, it is possible that ongoing gene flow may still be occurring in the contact zone. For example, work in the late 1960s by Smith (1969) suggested that cytotype hybrids occur in Louisiana and Mississippi, a finding that is at odds with our inference of no contemporary gene flow between the cytotypes. One way to reconcile these results is that early generation hybrids form but are not successful and do not contribute to on-going gene flow. The results of inter-cytotype crosses by Kasjaniuk et al. (2018) suggest strong F_1_ inviability in crosses between XYY males and XY females, but not in the other direction. This asymmetrical crossing result fits with our evidence for greater historical gene flow from XY into the XYY clade and with historical gene flow of a Y haplotype into the southern XYY clade. Further investigation of hybrid inviability beyond the F_1_, as well as the examination of gene flow in the contact zone should provide further insights into the scope for contemporary gene flow between the cytotypes.

### The large X-effect

Despite being at an early stage of divergence, our analyses suggest that the X chromosome contributes disproportionately to patterns of differentiation. Because we did not find comparable excess divergence on the neo-sex chromosomes or on the autosomes, the greater differentiation we observed on the ancestral sex chromosomes is likely due to divergence between cytotypes prior to the complete loss of gene flow at the second stage. Furthermore, we observed elevated differentiation even when using absolute measures of divergence, therefore differences in effective population size between the chromosomes alone are unlikely to explain our results. However, it is important to note that population genomic analysis cannot distinguish the relative importance of reduced gene flow from increased local adaptation in driving genetic divergence (Presgraves, 2018). In fact, these alternatives are not necessarily mutually exclusive. Our observations are consistent with a large body of work from animal systems from both crossing experiments and population genomic analysis that suggest greater reproductive isolation on the X chromosome (Coyne and Orr 2004; Presgraves 2018).

Our findings bolster similar patterns which have recently been observed in *Silene*, another plant system with heteromorphic sex chromosomes. In particular, studies based on both genomic scans (Hu and Filatov 2016) and genetic mapping of incompatibility loci (Brothers and Delph 2010; Demuth et al. 2014), have found a disproportionately large effect of the X chromosome on population differentiation and hybrid failure. In both *Rumex* and *Silene*, the observation of disproportionate differentiation on the X is unlikely to be completely explained by either a unmasking of deleterious (Muller 1940) or adaptive (Charlesworth et al. 1998) recessive alleles on the X, as gene loss on the Y is still in a relatively early stage. In our analyses, we observed excess X differentiation despite removing hemizygous genes. Nonetheless, it is possible that X-linked genes with retained Y-linked homologues still experience a greater effective dominance than autosomes due to a reduced expression of Y alleles. Given the evidence for widespread reduced Y homologue expression in *Rumex* (Hough et al. 2014), it is possible that the unmasking effect can act before gene loss from the Y chromosome.

There are a number of alternative explanations to the dominance theory that could also contribute to reduced gene flow on the X chromosome in *R. hastatulus*. As no signal of dosage compensation has been observed in *R. hastatulus* (Crowson et al. 2017), it is unlikely that this pattern can be explained by imbalances in dosage (Turelli and Orr 1995). Possible alternative explanations include a higher density of loci that confer incompatibility on the sex chromosomes (Masly and Presgraves 2007), evolution by genomic conflict, such as by meiotic drivers, which have been previously reported for *Rumex* (Wilby and Parker, 1988) or accelerated rates of pollen gene evolution on the sex chromosomes (Scott and Otto 2018; Sandler et al. 2018). Further analyses examining the heterogeneity in gene flow patterns across the X chromosome using both crossing experiments and population genomic analysis could help disentangle these possibilities further.

### The large Y-effect

We also found a disproportionate signal of population divergence on the Y chromosome of *R. hastatulus*, a pattern also reported in *S. latifolia* (Muir et al. 2011). While we can attribute part of the signal involving high values of *F*_ST_ to linked selection, in contrast to *S. latifolia*, values of absolute divergence are also disproportionately high on the Y chromosome and importantly even higher than those on the X. This suggests that, like the X chromosome, the Y chromosome in *R. hastatulus* contributes disproportionately to reproductive isolation and/or local adaptation, a so-called ’large Y effect’. Given the general paucity of retained Y genes on animal Y chromosomes, tests for a large Y effect are near absent. To our knowledge our study is the first to observe this effect after controlling for differences in effective population size. Although the Y chromosome is typically highly degenerate and would not be expected to contribute disproportionately to reproductive isolation and adaptation, Y-specific adaptation to pollen competition and male reproduction (Sandler et al 2017) on a relatively young Y chromosome could be an important driver of reproductive isolation. If, as predicted by theory (Scott and Otto 2018), haploid selection during pollen competition is elevated on the Y chromosome, this could contribute to a dominance effect on the Y driving an accumulation of factors on this chromosome causing reproductive isolation. One possible explanation for the absence of a large-Y effect in *S. latifolia* is that population differentiation may be more limited than we observe between the cytotypes of *R. hastatulus.* Reduced gene flow, local adaptation and an accumulation of factors contributing to reproductive isolation on the Y may have allowed for higher level of genetic divergence in our study system.

### Neo-sex chromosomes and chromosome turnover

Our findings suggest that sex chromosome turnover may have contributed to the cessation of gene flow between *R. hastatulus* cytotypes. Although the loss of recombination between the neo-X and neo-Y is estimated to be older than the loss of gene flow between cytotypes, our estimate of the time of neo-sex chromosome formation represents the age of initial loss of recombination between the neo-sex chromosomes and is therefore a conservative estimate for the time since fixation of the rearrangement in the XYY geographical range. Similar values of *d*_A_ between autosomal and neo-sex-linked loci suggest that both types of genes have been diverging at similar rates during the same amount of time. This observation is consistent with the possibility that sex chromosome turnover or neo-sex chromosome formation caused the loss of gene flow between the cytotypes. However, this pattern could also arise if the origin of the neo-sex chromosomes and the loss of gene flow were coincident in time, without a causal role. This conundrum highlights a shortcoming of genomic studies of genetic divergence: if a locus contributes to complete isolation between populations, the signal of its contribution to divergence will be indistinguishable from the general loss of gene flow across the genome. Nonetheless, the similar timescales we estimated for these events combined with evidence from crossing studies that demonstrate the important effect of the rearrangement on cytotype hybrid fitness (Kasjaniuk et al. 2018) suggest a causal relation between sex chromosome turnover, genetic divergence and reproductive isolation between the cytotypes of *Rumex hastatulus.*

### Data archiving

Nucleotide sequences are stored in the NCBI SRA databank, under BioProject PRJNA522871.

### Author contributions

S.I.W., S.C.H.B., and F.E.G.B. conceived of and designed the study, as well contributed to writing of the paper. F.E.G.B. collected and analyzed the data.

## Acknowledgements

The authors would like to thank Tia Harrison for help with DNA and RNA extraction, Wei Wang for coding scripts, and Yunchen Gong for server help. This research was funded by Discoveries grants from the Natural Sciences and Engineering Research Council of Canada to SCHB and SIW. In memory of Josh Hough, whose intellectual curiosity and passion for research launched our investigations into *Rumex* sex chromosome evolution.

