## Supporting Information for "Ancestral and neo-sex chromosomes contribute to population divergence in a dioecious plant"

**Supplementary Table 1.** Population sampling and sequencing information for *Rumex hastatulus*. Y-haplotype information based on ML tree. Demographic and geographic information in Pickup and Barrett (2013)

| Population | GBS samples | RNA samples | Y-haplotype group |
|---|---|---|---|
| AL-BRE | 2 | 1 | NC |
| AL-BRU | 4 | 1 | NC |
| FL-JAS | 4 | 1 | FL |
| FL-MAR | 4 | 1 | *(Y SNPs not found)* |
| FL-HAM | 0 | 2 | FL |
| GA-BEL | 3 | 3 | FL |
| GA-GLA | 6 | 3 | FL |
| GA-STA | 6 | 3 | Mix: NC + FL |
| LA-BEN | 3 | 1 | TX |
| LA-DER | 0 | 2 | TX |
| NC-BAT | 3 | 1 | NC |
| NC-ELI | 6 | 1 | *(Y SNPs not found)* |
| NC-HIC | 0 | 1 | *(Y SNPs not found)* |
| NC-KIN | 0 | 1 | NC |
| NC-ROS | 4 | 1 | NC |
| OK-BAC | 6 | 1 | TX |
| OK-RAT | 6 | 3 | TX |
| OK-WIL | 0 | 2 | TX |
| SC-BRA | 5 | 3 | NC |
| SC-MAR | 6 | 1 | NC |
| SC-PRO | 7 | 3 | Mix: NC + FL |
| TX-ATH | 5 | 3 | TX |
| TX-LIV | 4 | 3 | TX |
| TX-MTP | 1 | 3 | TX |

| | | | |
|---|---|---|---|
| TX-OAK | 2 | 1 | TX |
| TX-ROS | 5 | 1 | TX |
| **TOTAL** | **92** | **47** | **21** |

**Supplementary Table 2.** Best fit parameters from *δaδi* model as output from dadi, *ie* in units of *N*e, and scaled by our estimate of ancestral population size in *Rumex hastatulus*.

| | Including Alabama | | Excluding Alabama | |
|---|---|---|---|---|
| | Relative | Scaled ($10^3$) | Relative | Scaled ($10^3$) |
| Ancestral Ne | | 316 | | 150 |
| T1 | 3.4 | 2,149 | 5.90 | 1,250 |
| N1 XY | 2.90 | 916 | 5.92 | 627 |
| N1 XYY | 0.61 | 193 | 1.55 | 164 |
| XY → XYY | 2.00 | 1,264 | 0.87 | 184 |
| XYY → XY | 0.12 | 76 | 0.10 | 21 |
| T2 | 0.079 | 50 | 0.22 | 66 |
| N2 XY | 4.59 | 1,450 | 15.23 | 1,614 |
| N2 XYY | 11.90 | 3,760 | 7.32 | 742 |

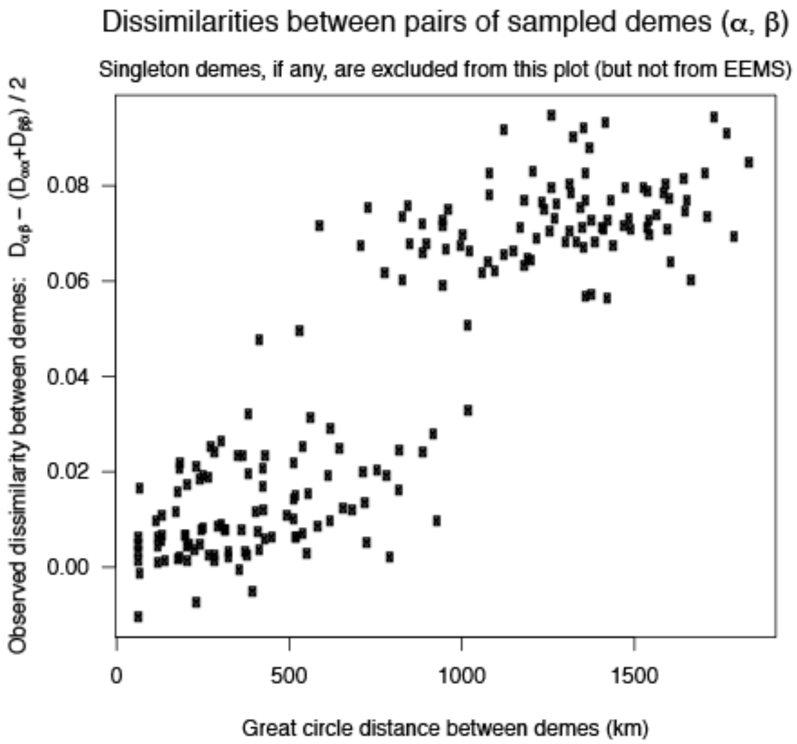


**Supplementary Figure 1.** EEMS estimate of dissimilarities between pairs of sampled demes of *Rumex hastaulus*

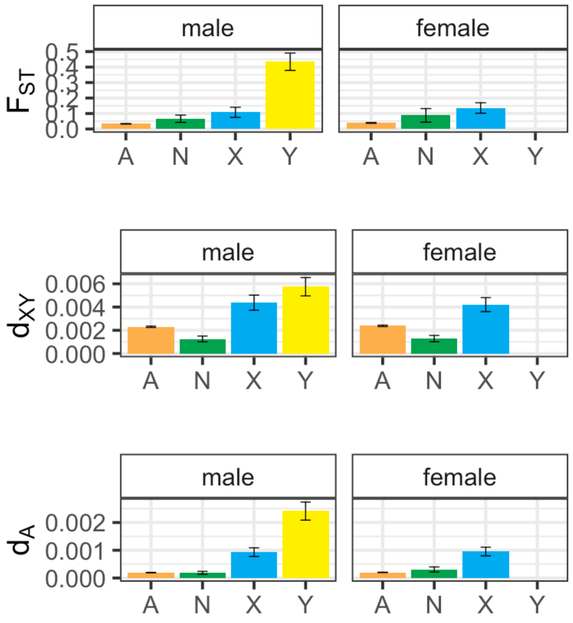


**Supplementary Figure 2.** Male to female sets for values of divergence in *Rumex hastatulus*

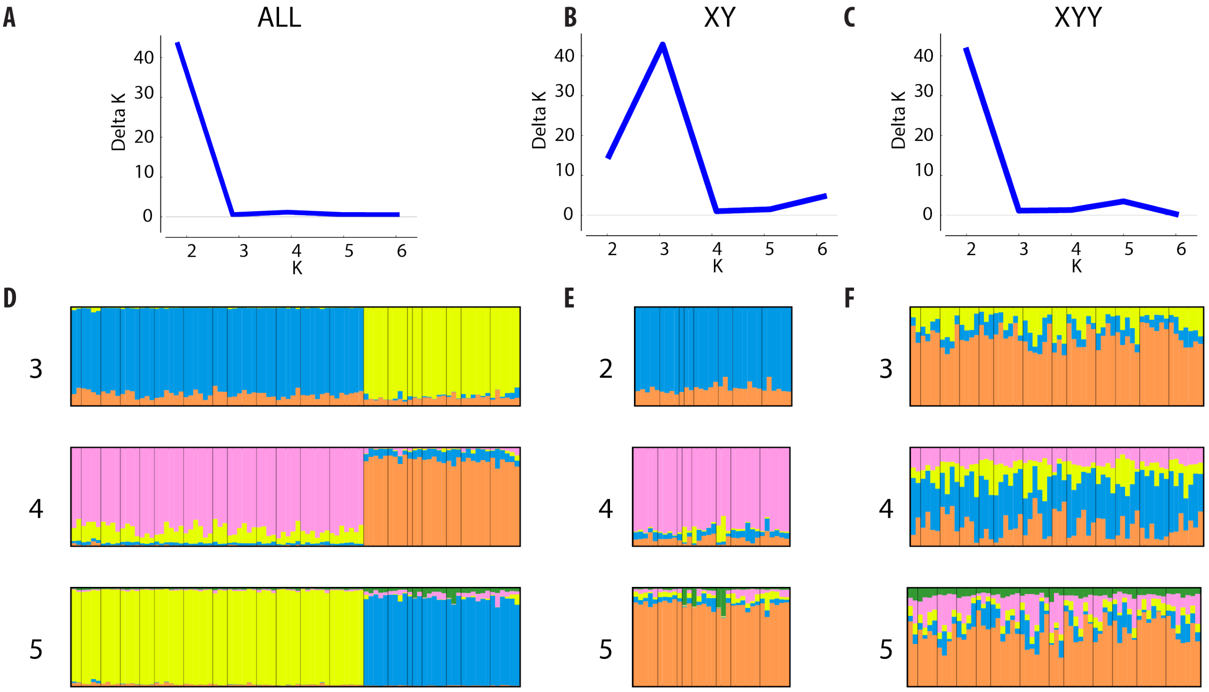


**Supplementary Figure 3. A-C** Likelihood distribution for the number of clusters for populations of *Rumex hastatulus* represented in the GBS SNP from StructureHarvester. **D-F** STRUCTURE plot for less likely values of K for each cytotype.

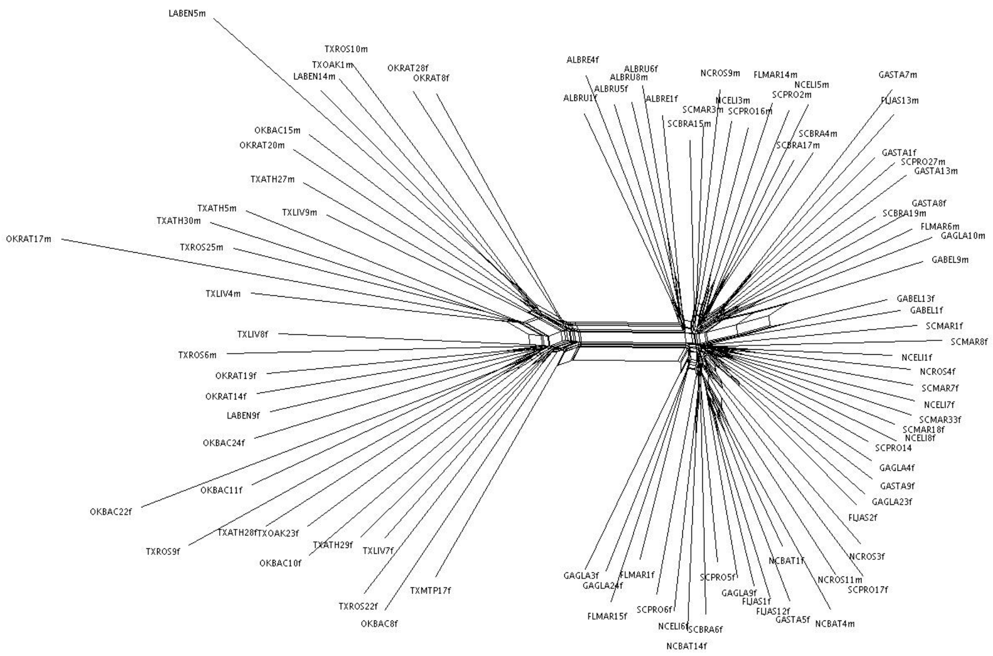


**Supplementary Figure 4.** Splitstree for SNPs from the GBS dataset based on population samples *of Rumex hastatulus* from throughout its range.

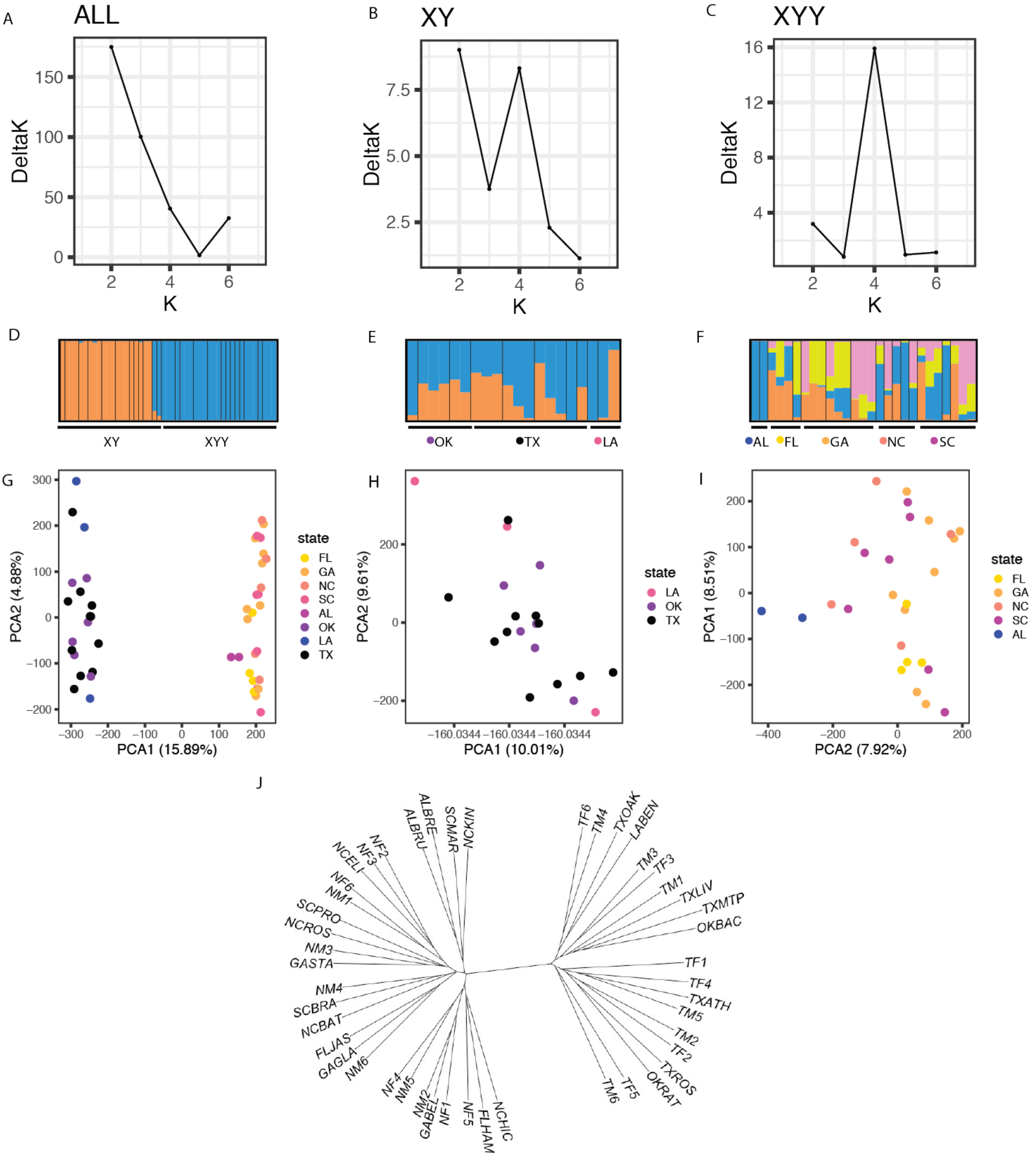


**Supplementary Figure 5.** Evidence of divergence between populations of the two cytotypes of *Rumex hastatulus* using SNP from RNAseq data. **A-C** most likely number of clades as estimated from the Evanno method in StructureHarvester **D-F** Structure plot for each cytotype, **G-I** Principal component analysis (PCA) for each cytotype, **J** neighbor-joining tree.

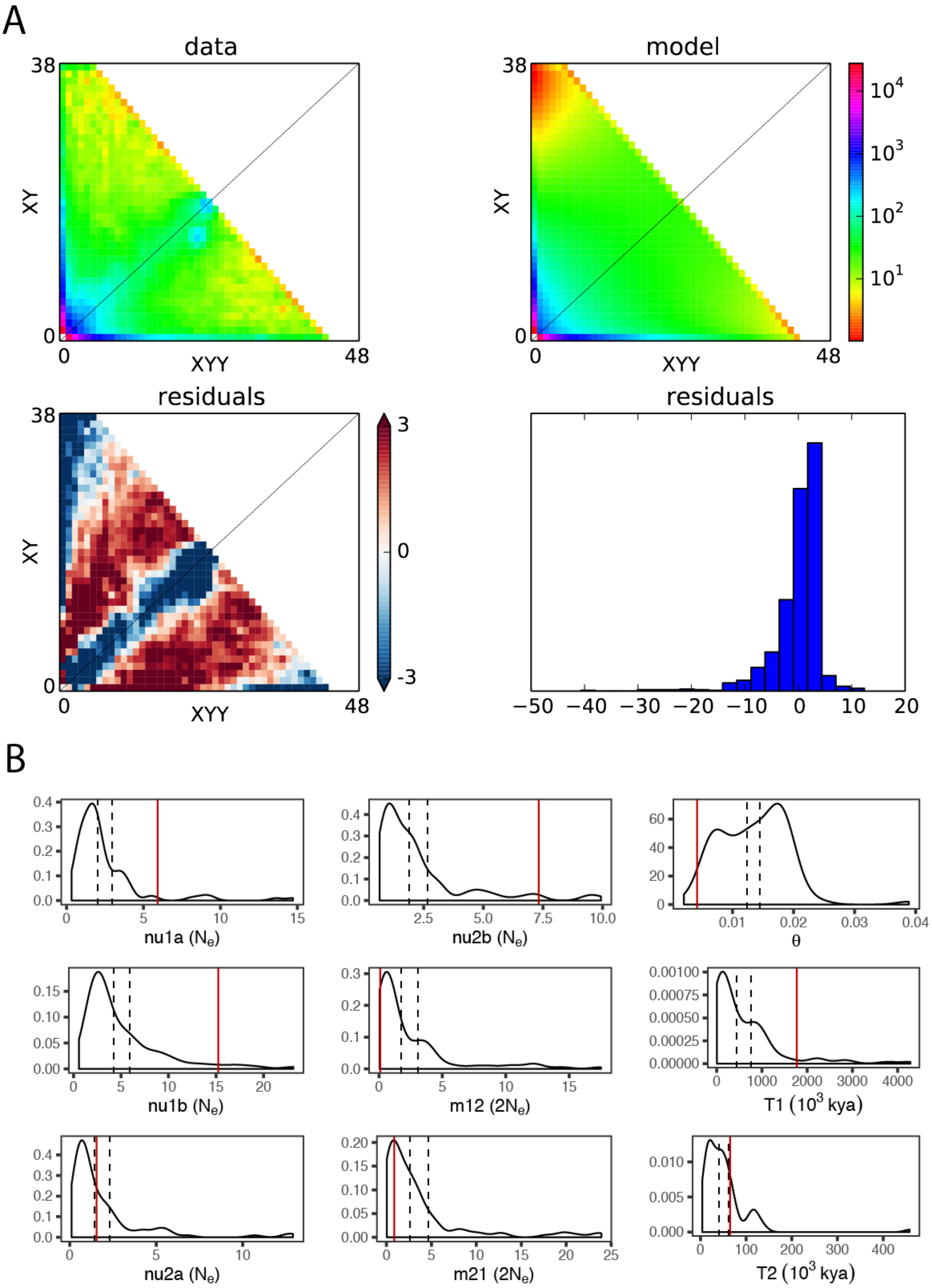


**Supplemental Figure 6.** *Rumex hastatulus* demographic model fitting from *δaδi* **A** residuals and **B** confidence intervals from bootstrapping

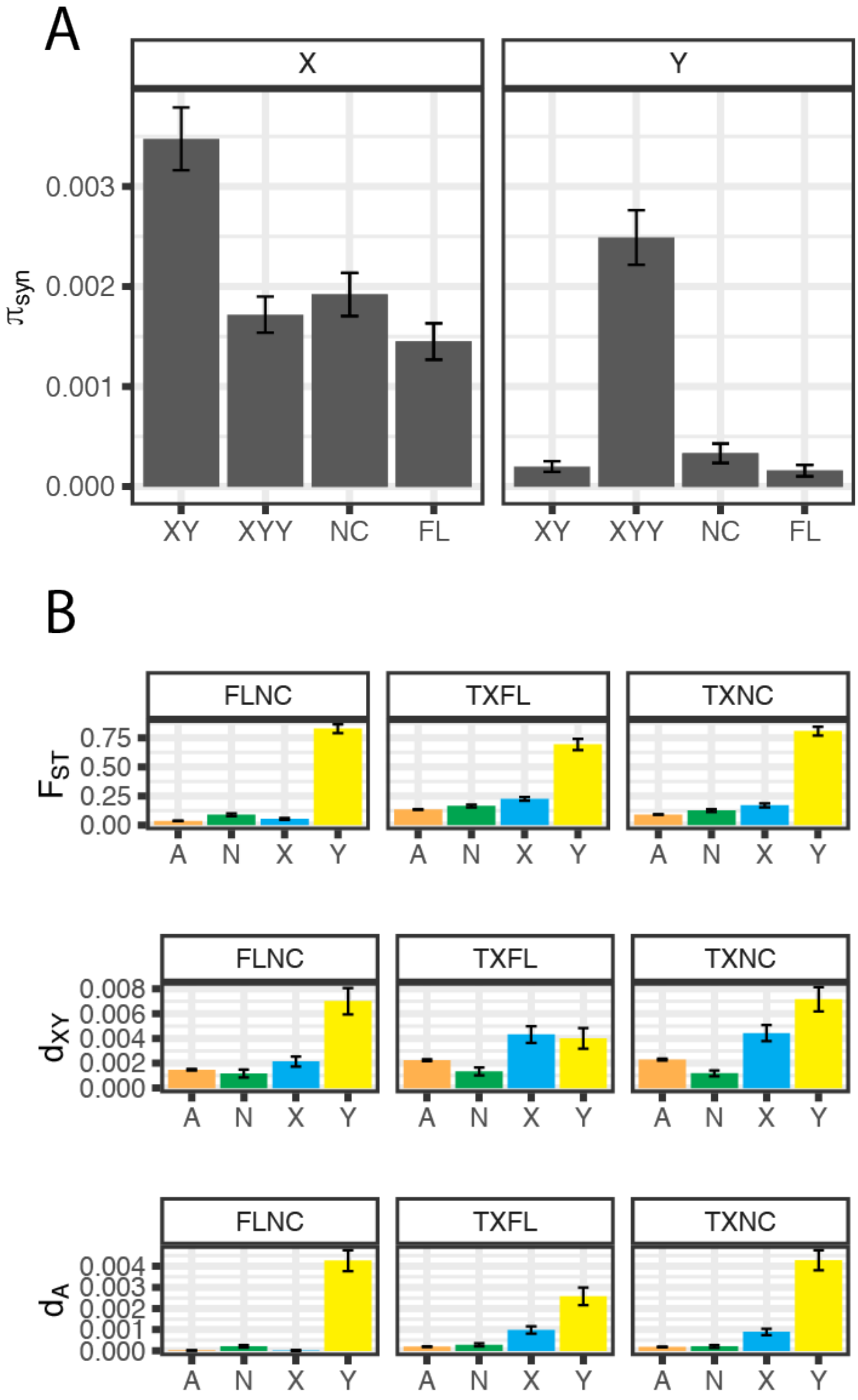


**Supplementary Figure 7. A.** $\pi_{syn}$ at X and Y loci in both cytotypes, and the Y haplotypes subclades of *Rumex hastatulus*, and **B.** $F_{ST}$, $d_{XY}$ , and $d_A$ for each Y haplotype subclade

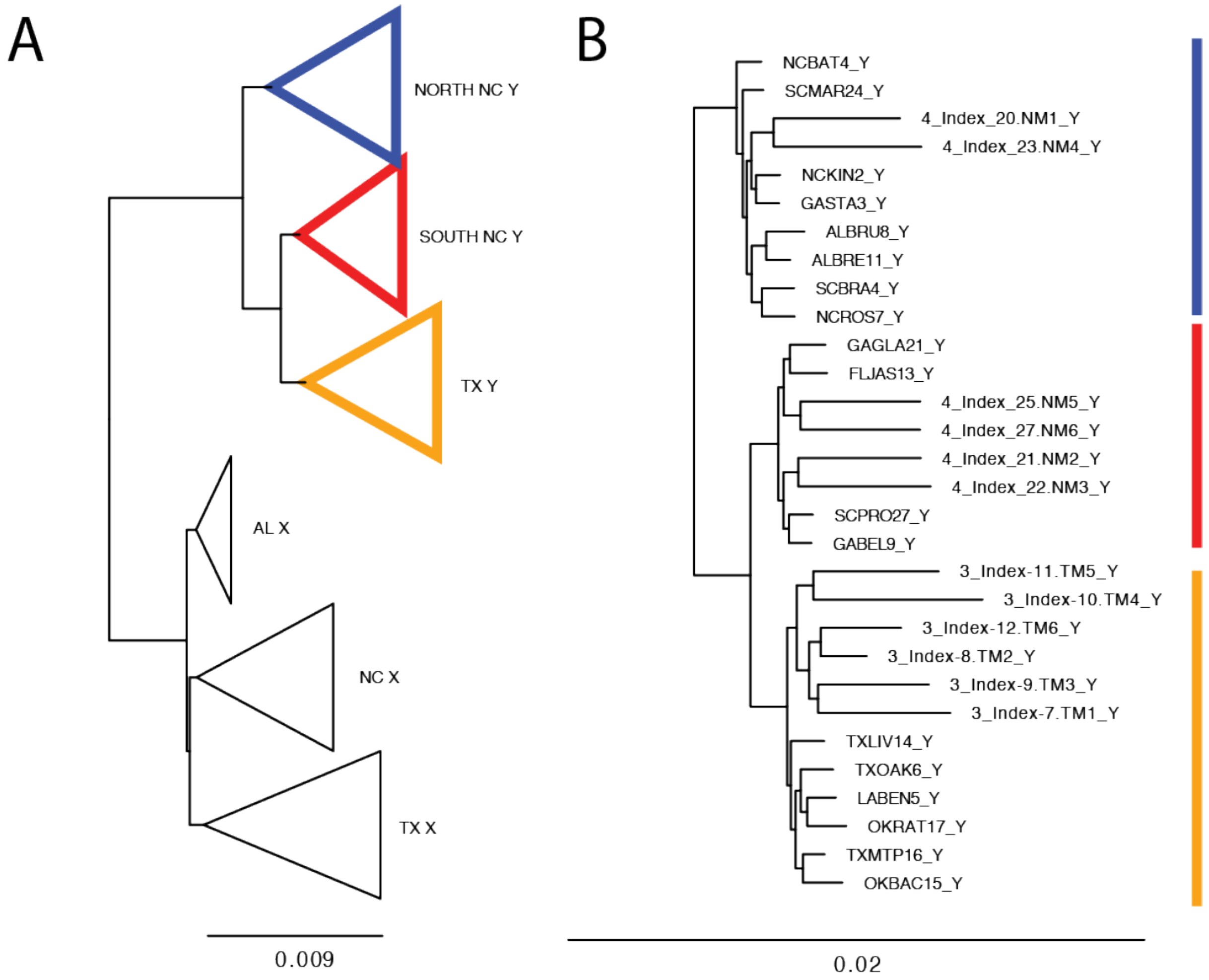


**Supplementary Figure 8.** Phased ancestral sex chromosome haplotype tree for **A** all haplotypes, and **B** only Y haplotypes of *Rumex hastatulus*.
